# The Preclinical Inhibitor GS441524 in Combination with GC376 Efficaciously Inhibited the Proliferation of SARS-CoV-2 in the Mouse Respiratory Tract

**DOI:** 10.1101/2020.11.12.380931

**Authors:** Yuejun Shi, Lei Shuai, Zhiyuan Wen, Chong Wang, Yuanyuan Yan, Zhe Jiao, Fenglin Guo, Zhen F. Fu, Huanchun Chen, Zhigao Bu, Guiqing Peng

**Author notes:** These authors contributed equally to this work (Yuejun Shi, Lei Shuai). Correspondence to Zhigao Bu and Guiqing Peng.

## Abstract

The unprecedented coronavirus disease 2019 (COVID-19) pandemic, caused by severe acute respiratory syndrome coronavirus 2 (SARS-CoV-2), is a serious threat to global public health. Development of effective therapies against SARS-CoV-2 is urgently needed. Here, we evaluated the antiviral activity of a remdesivir parent nucleotide analog, GS441524, which targets the coronavirus RNA-dependent RNA polymerase enzyme, and a feline coronavirus prodrug, GC376, which targets its main protease, using a mouse-adapted SARS-CoV-2 infected mouse model. Our results showed that GS441524 effectively blocked the proliferation of SARS-CoV-2 in the mouse upper and lower respiratory tracts via combined intranasal (i.n.) and intramuscular (i.m.) treatment. However, the ability of high-dose GC376 (i.m. or i.n. and i.m.) was weaker than GS441524. Notably, low-dose combined application of GS441524 with GC376 could effectively protect mice against SARS-CoV-2 infection via i.n. or i.n. and i.m. treatment. Moreover, we found that the pharmacokinetic properties of GS441524 is better than GC376, and combined application of GC376 and GS441524 had a synergistic effect. Our findings support the further evaluation of the combined application of GC376 and GS441524 in future clinical studies.

**Importance:** Severe acute respiratory syndrome coronavirus 2 (SARS-CoV-2) causes coronavirus disease 2019 (COVID-19), which has seriously threatened global public health and economic development. Currently, effective therapies to treat COVID-19 are urgently needed. In this study, we assessed the efficacy of the preclinical inhibitors GC376 and GS441524 using a mouse-adapted SARS-CoV-2 infected mouse model for the first time. Our results showed that low-dose combined application of GC376 and GS441524 could effectively protect mice from HRB26M infection in the upper and lower respiratory tracts. Hence, the combined application should be developed and considered for future clinic practice.

## Introduction

Severe acute respiratory syndrome coronavirus 2 (SARS-CoV-2) causes coronavirus disease 2019 (COVID-19), which spread rapidly to more than 235 countries. The number of infections has exceeded 40 million, and more than 1 million deaths were reported (https://www.who.int/emergencies/diseases/novel-coronavirus-2019). The World Health Organization (WHO) declared COVID-19 a global health emergency. Multiple vaccine candidates and therapeutics are in clinical trials (1–6), and effective treatments or cures for COVID-19 are still urgently needed.

Coronaviruses are enveloped, positive-sense, single-stranded RNA viruses, and their genomic RNA is approximately 30 kb and contains at least 6 open reading frames (ORFs) (7). The first ORF (ORF 1a/b) encodes two polyproteins, pp1a and pp1ab, and these polyproteins are processed via a main protease (M^pro^, also known as the 3C-like protease) and one or two papain-like proteases (PLPs) into 16 nonstructural proteins (nsps) (8, 9). These nsps engage in the production of subgenomic RNAs that encode four main structural proteins [envelope (E), membrane (M), spike (S), and nucleocapsid (N) proteins] and other accessory proteins (8, 9). Among these nsps, M^pro^ and nsp12 (RdRp) of SARS-CoV-2 are functionally and structurally conserved among these viruses and essential for viral replication; they are considered a potential target for the design of antiviral drugs (10–16).

GC376, a dipeptidyl bisulfite adduct salt, exerts strong inhibitory effects on picornaviruses and coronaviruses (11, 14, 15, 17). The latest research has shown that GC376 can also inhibit the replication of SARS-CoV-2 in Vero E6 cells (12, 13, 16). Moreover, antiviral treatment with GC376 led to a full recovery in laboratory cats with feline infectious peritonitis (FIP) caused by feline coronavirus (FIPV) (18). Currently, no studies have evaluated the ability of GC376 to inhibit SARS-CoV-2 in animals. In addition, remdesivir (RDV) and its parent nucleoside analog GS-441524 inhibit CoVs and other viruses (19–22). The latest research also suggests that RDV inhibits SARS-CoV-2 *in vivo* and *in vitro* (10, 23–27). However, recent research showed that RDV was not intended for lung-specific delivery, and GS-441524 is the predominant metabolite that reaches the lungs, which suggests that GS-441524 is superior to RDV for COVID-19 treatment (28, 29).

Because GC376 and GS441524 target the key proteases M^pro^ and RdRp of SARS-CoV-2 replication, respectively, their combined application may be more effective in the treatment of COVID-19. The combined application of GC376 with RDV produced additive sterilizing additive effects against SARS-CoV-2 in Vero E6 cells (13). However, the efficacy of the combined application of GC376 and GS441524 in the inhibition of SARS-CoV-2 replication was not investigated *in vivo* and *in vitro*. Our team generated a mouse-adapted SARS-CoV-2 (HRB26M) that efficiently replicates in the upper and lower respiratory tract in BALB/c mice (4, 30). Hence, we assessed the antiviral efficacy of GC376 and GS441524 in an HRB26M-infected BALB/c mouse model. Our results showed that GC376, which effectively inhibited the virus at the cellular level, was less effective *in vivo*. The ability of GS441524 to block the proliferation of SARS-CoV-2 was significantly higher than GC376 in mice. Notably, the low-dose combined application of GC376 and GS441524 could effectively block SARS-CoV-2 infection in the mouse upper and lower respiratory tract.

## Results and Discussion

### Structural analysis of GC376 and GS441524 targeting SARS-CoV-2 M^pro^ and nsp12 (RdRp)

In this study, we first analyzed the mechanism by which GC376 and GS441524 inhibit SARS-CoV-2 replication through structural biology methods. GC376 (**Fig. 1a**) is a preclinical inhibitor against feline infectious peritonitis virus (FIPV) (31) that has become a broad-spectrum prodrug targeting coronavirus M^pro^ (14, 32, 33). We determined the high-resolution crystal complex structure of M^pro^ with GC376 at a resolution of 2.35 Å (**Table S1**). The crystal of M^pro^-GC376 belongs to the space group P1211, and an asymmetric unit contains two molecules (designated protomer A and protomer B) (**Fig. 1b**). The structure of each protomer contains three domains with the substrate-binding site comprised of a His41-Cys145 dyad located in the cleft between domain I and II (**Supplementary Fig. 1**). The electron density map clearly showed compound GC376 in the substrate binding pocket of the SARS-CoV-2 M^pro^, and details of the interaction are shown in **Fig. 1c and Supplementary Fig. 1**. GC376 interacts with His41, Phe140, Leu141, Asn142, Ser144, Cys145, His163, His164, Met165, Glu166, His172, Asp187, Arg188 and Gln189 through many hydrogen bonds and hydrophobic interactions, and binds stably in the groove formed by domain I and domain II. Moreover, we found that GC376 is covalently attached to Cys145 as a hemithioacetal (**Fig. 1c**), which prevents the binding and processing of M^pro^ to the substrate (12, 16, 32). Additionally, GS441524 is the parent nucleoside of RDV (**Fig. 1d and Supplementary Fig. 2a**), and has shown broad-spectrum inhibition of the replication of various coronaviruses, including SARS-CoV-2 (28, 34, 35). Since the mechanism by which GS441524 inhibits SARS-CoV-2 replication is similar to that of RDV (28, 34), we showed the cryo-electron microscopy (cryo-EM) ternary structure of the complex with RDV (PDB: 7BV2) (10). Previous studies have shown that RDV, like many nucleotide analog prodrugs, inhibits viral RdRp activity through nonobligate RNA chain termination (10, 36, 37). Moreover, we observed that RDV stably bound nsp12 through hydrophobic interactions with Arg555, Asp623, Ser682, Thr687 and Asp760 (**Fig. 1e, f and Supplementary Fig. 2b**).

**Fig.1.**
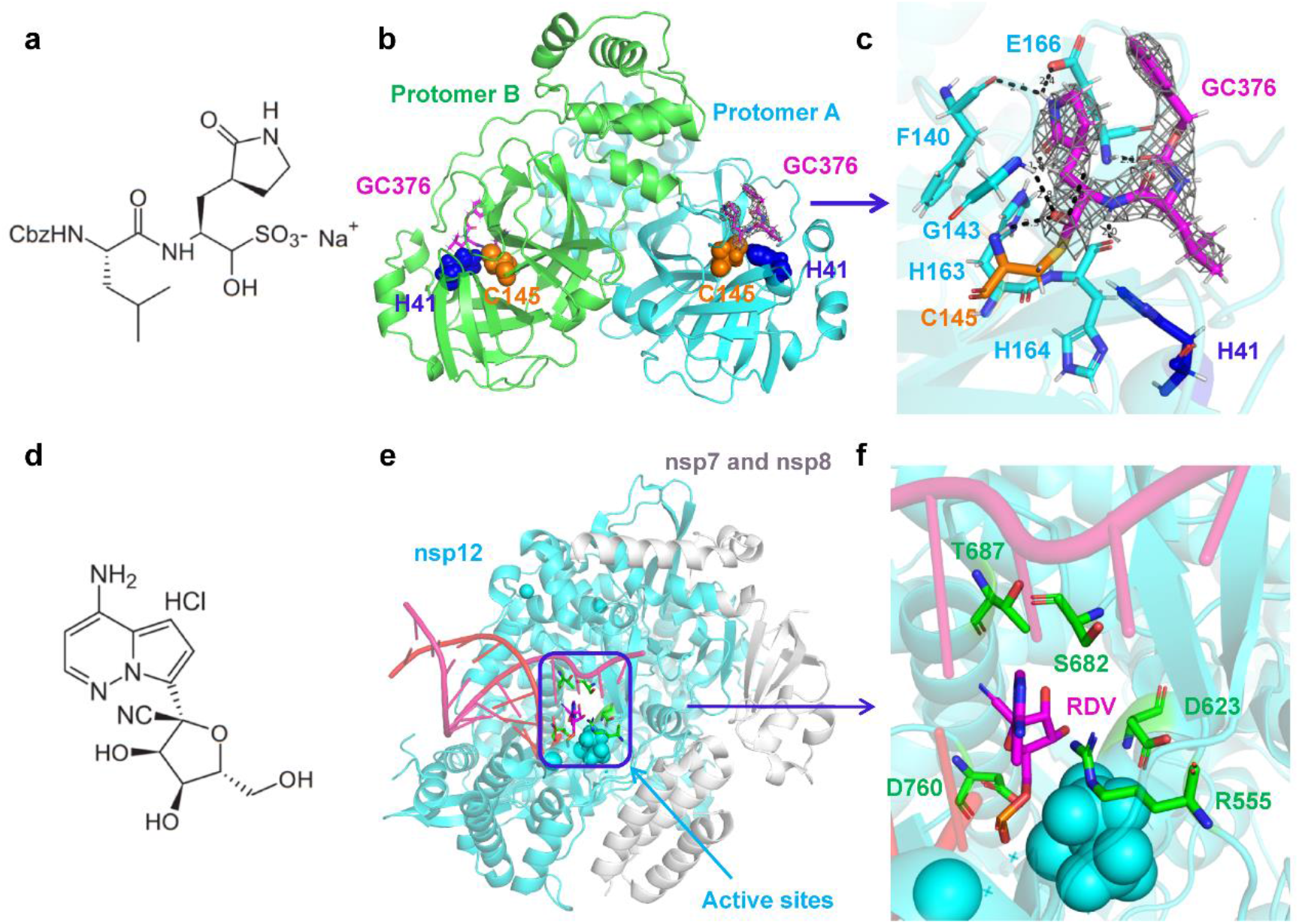
Structural analysis of GC-376 and GS441524 targeting SARS-CoV-2 M^pro^ and RdRp. (a) The dipeptidyl protease inhibitor, GC376. (b) Crystal structure of SARS-CoV-2 M^pro^ in complex with GC376. (c) GC376 interacts covalently with the active cysteine site of SARS-CoV-2 M^pro^. Electron density at 1.5 σ is shown in gray mesh. Hydrogen bonds are shown as red dashed lines. (d) The chemical structure of GS-441524. (e) Cryo-EM structure of the apo nsp12-nsp-7-nsp8 RdRp complex (PDB ID: 7BV2). (f) Enlarged view of the active site, depicting the interaction between RDV and surrounding amino acids.

### Combined application of GC376 and GS441524 has an enhanced ability to inhibit SARS-CoV-2 in Vero E6 cells

We evaluated the inhibitory efficacy of GC376, GS441524 and the combined application of GC376 and GS441524 (molar ratio: 1:1) (GC376+GS441524) on the replication of live virus (SARS-CoV-2: HRB26 and HRB26M) in Vero E6 cells. We first tested the cellular cytotoxicity of these compounds *in vitro*. GC376 and GS441524 did not produce obvious cytotoxicity at concentrations up to 250 μM in Vero E6 cells (CC_50_ > 250 μM**; Supplementary Fig. 3**). Our results showed that GC376 and GS441524 were efficacious against HRB26, with 50% inhibitory concentration (EC_50_) values of 0.643±0.085 μM and 5.188±2.476 μM, respectively (**Fig. 2a, b**). GC376 and GS441524 were also efficacious against HRB26M, with EC_50_ values of 0.881±0.109 μM and 5.047±2.116 μM, respectively (**Fig. 2d, e**). Our results showed that the ability of GC376 to inhibit viral (HRB26 and HRB26M) replication was better than GS441524 when the agents were applied alone. We observed that the GC376+GS441524 more effectively inhibited HRB26 and HRB26M replication than single treatment, with EC_50_ values of 0.531±0.162 μM and 0.369±0.068 μM, respectively (**Fig. 2c, f**). In the replication process of coronavirus, M^pro^ is one of the first nsps processed by the polyprotein, and other replication-related proteins, such as RdRp, can be produced with the participation of M^pro^ and PLP proteases (8, 9). This phenomenon may be the reason why GC376 is better than GS441524, and their combined application may result in a synergistic effect because these agents target different proteins involved in virus replication.

**Fig.2.**
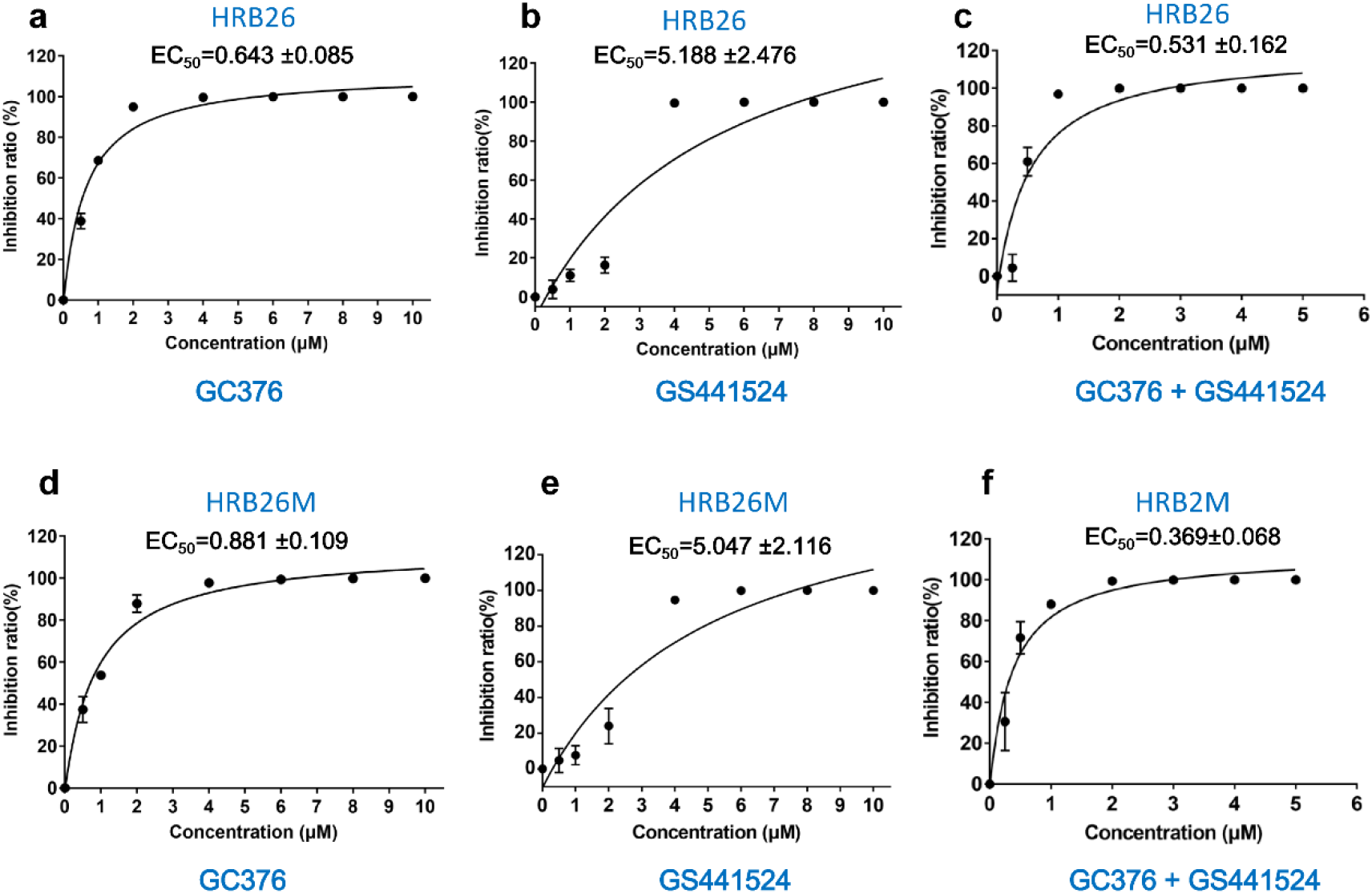
GC376 and GS441524 potently inhibit SARS-CoV-2 in Vero E6 cells. (a-c) Percent inhibition of SARS-CoV-2 (HRB26) by GC376, GS441524 and GC376+GS441524 in Vero E6 cells. (d-f) Percent inhibition of SARS-CoV-2 (HRB26M) by GC376, GS441524 and GC376+GS441524 in Vero E6 cells. The concentrations of GC376 and GS441524 ranged from 0 to 10 μM. The 50% inhibitory concentration (EC_50_) was calculated using GraphPad Prism 7.0 to assess inhibition ratios at different inhibitor concentrations. The error bars show the S.D. of the results from three replicates.

### Low-dose GC376+GS441524 could effectively protect mice from HRB26M infection in the upper and lower respiratory tracts

The potential toxicity of GC376, GS441524 and GC376+GS441524 were evaluated in healthy specific pathogen-free (SPF) mice at 4-6 weeks. The mice were intramuscularly (i.m.) administered GC376 (40 mM/l, 100 μl), GS441524 (40 mM/l, 100 μl) and GC376+GS441524 (20 mM/l, 100 μl) for 15 days, and they were observed daily for adverse effects. During the study period, there were no clinically significant changes in vital signs or clinical laboratory parameters (**Supplementary Fig. 4**), indicating that the dose and administration route of GC376, GS441524 and GC376+GS441524 were suitable for the *in vivo* efficacy study.

We assessed the antiviral efficacy of i.m. administered GC376 (40 or 8 mM/l, 100 μl), GS441524 (40 or 8 mM/l, 100 μl) and GC376+GS441524 (20 or 4 mM/l, 100 μl) against SARS-CoV-2 in mice. In the compound-treated group, the viral RNA and viral titer were detected in the nasal turbinates and lungs on days 3 and 5 post-inoculation (p.i.); the viral RNA loads and viral titer in the nasal turbinates were not significantly decreased compared with those in the mock-treated mice on day 3 p.i (**Fig. 3a, b and Supplementary Fig. 5a, b**). Although no significant decreases in viral RNA loads were observed, high-dose GC376+GS441524 significantly inhibited the replication of SARS-CoV-2 in the turbinate on day 5 p.i. (**Fig. 3c, d and Supplementary Fig. 5c, d**). Our results showed that high-dose GS441524 and GC376+GS441524 completely inhibited the proliferation of SARS-CoV-2 in mouse lungs on days 3 and 5 p.i. (**Fig. 3e f, g, h and Supplementary Fig. 6**). Besides, the virus yields in the lungs of the high-dose GC376-treated group were significantly reduced compared to mock-treated mice on day 5 p.i., but it could not completely block the viral proliferation (**Fig. 3h and Supplementary Fig. 6d**). Additionally, the ability of the low-dose compound-treated group to inhibit viral replication was lower than the high-dose group in the lungs (**Fig. 3e, f, g, h and Supplementary Fig. 6**). Similar to RDV (30), high-dose GS441524 and GC376+GS441524 completely blocked the proliferation of SARS-CoV-2 in mouse lungs, but these agents failed to block virus proliferation in the upper respiratory turbinate on day 5 p.i. These data indicated that high-dose i.m. compound treatment (excluding GC376) efficiently inhibited the replication of SARS-CoV-2 in the lungs, but not in the nasal turbinates of BALB/c mice.

**Fig.3.**
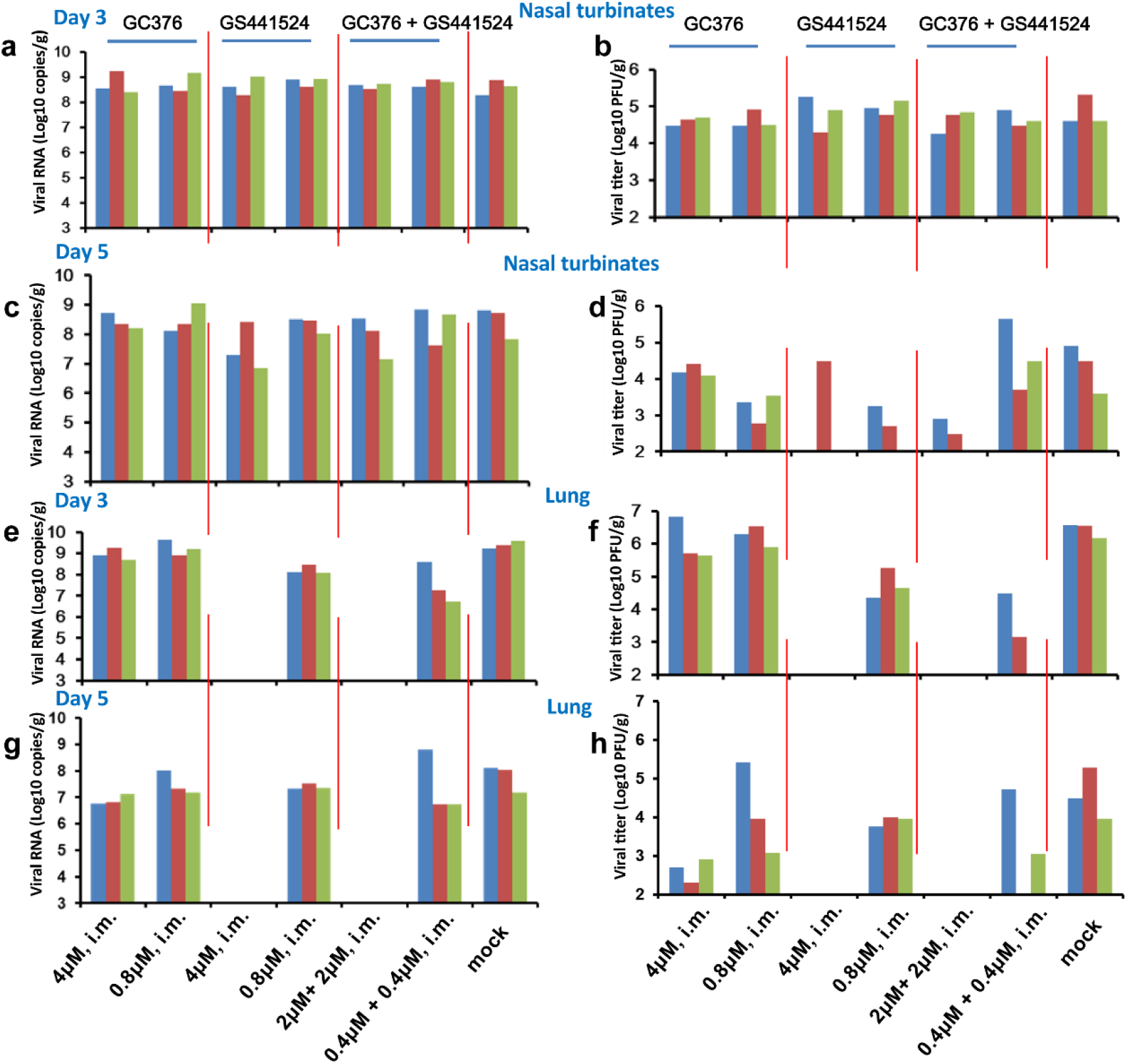
Evaluation of i.m. GC376, GS441524 and GC376+GS441524 against SARS-CoV-2 infection in mice. Four- to six-week-old female BALB/c mice were intramuscularly administered a loading dose of GC376 (4 or 0.8 μM), GS441524 (4 or 0.8 μM), GC376+GS441524 (2 μM+2 μM or 0.4 μM+0.4 μM), followed by a corresponding daily maintenance dose. Control mice were administered vehicle solution (12% sulfobutylether-β- cyclodextrin, pH 3.5) daily, in parallel (0 μM). One hour after administration of the loading dose of GC376, GS441524, GC376+GS441524 or vehicle solution, the mice were inoculated intranasally with 10^3.6^ PFU of HRB26M in a volume of 50 μl. On days 3 and 5 p.i., three mice in each group were euthanized and their nasal turbinates and lungs were collected. The viral RNA copies and infectious titers in the nasal turbinates (a-d) and lungs (e-h) were detected by qPCR and viral titration. The concentrations of the daily maintenance doses are shown.

Furthermore, we assessed whether intranasal (i.n.) administration would improve the efficacy of GC376 and GS441524 to inhibit the replication of SARS-CoV-2 in the upper respiratory tract. Mice were treated with GC376 (i.n. and i.n.+i.m.), GS441524 (i.n. and i.n.+i.m.) and GC376+GS441524 (i.n. and i.n.+i.m.). Our results showed that GS441524 and GC376+GS441524 markedly inhibited viral replication in nasal turbinates on days 3 and 5 p.i. (**Fig. 4b, d and Supplementary Fig. 7b, d**). Only slightly infectious virus was detected on day 5 p.i., although viral RNA was detected in 2 of the 3 mice in the GS441524-treated group (**Fig. 4c, d and Supplementary Fig. 7c, d**). However, i.n. administration of GS441524 did not significantly inhibit viral replication in the lungs compared to that of the mock-treated group on day 5 p.i (**Fig. 4g, h and Supplementary Fig. 8c, d**). Notably, we observed that i.n. administration of GC376+GS441524 blocked viral replication in the lungs on day 5 p.i., although viral RNA was detected in 1 of the 3 mice (**Fig. 4 g, h and Supplementary Fig. 8 c, d**). GS441524 treatment via i.n. and i.m. routes completely blocked viral replication in the nasal turbinates and lungs, although viral RNAs were detected from the nasal turbinates of 2 of the 3 mice on day 3 p.i (reduced approximately 700-fold compared to that of the mock-treated group) (**Fig. 4 and Supplementary Figs. 7 and 8**). Similarly, low-dose GC376+GS441524 treatment (i.n.+ i.m.) also completely blocked viral replication in the nasal turbinates and lungs, although viral RNAs were detected from the nasal turbinates of 2 of the 3 mice on day 3 p.i. and 5 p.i. (reduced approximately 1500-fold and 263-fold compared to that of the mock-treated group, respectively) (**Fig. 4 and Supplementary Figs. 7 and 8**). Although GC376 treatment via the i.n. or combined i.n. and i.m. routes reduced the level of virus replication on day 5 p.i **(Supplementary Figs. 7d and 8d**), it did not completely blocked viral replication in nasal turbinates and lungs (**Fig. 4**).

**Fig.4.**
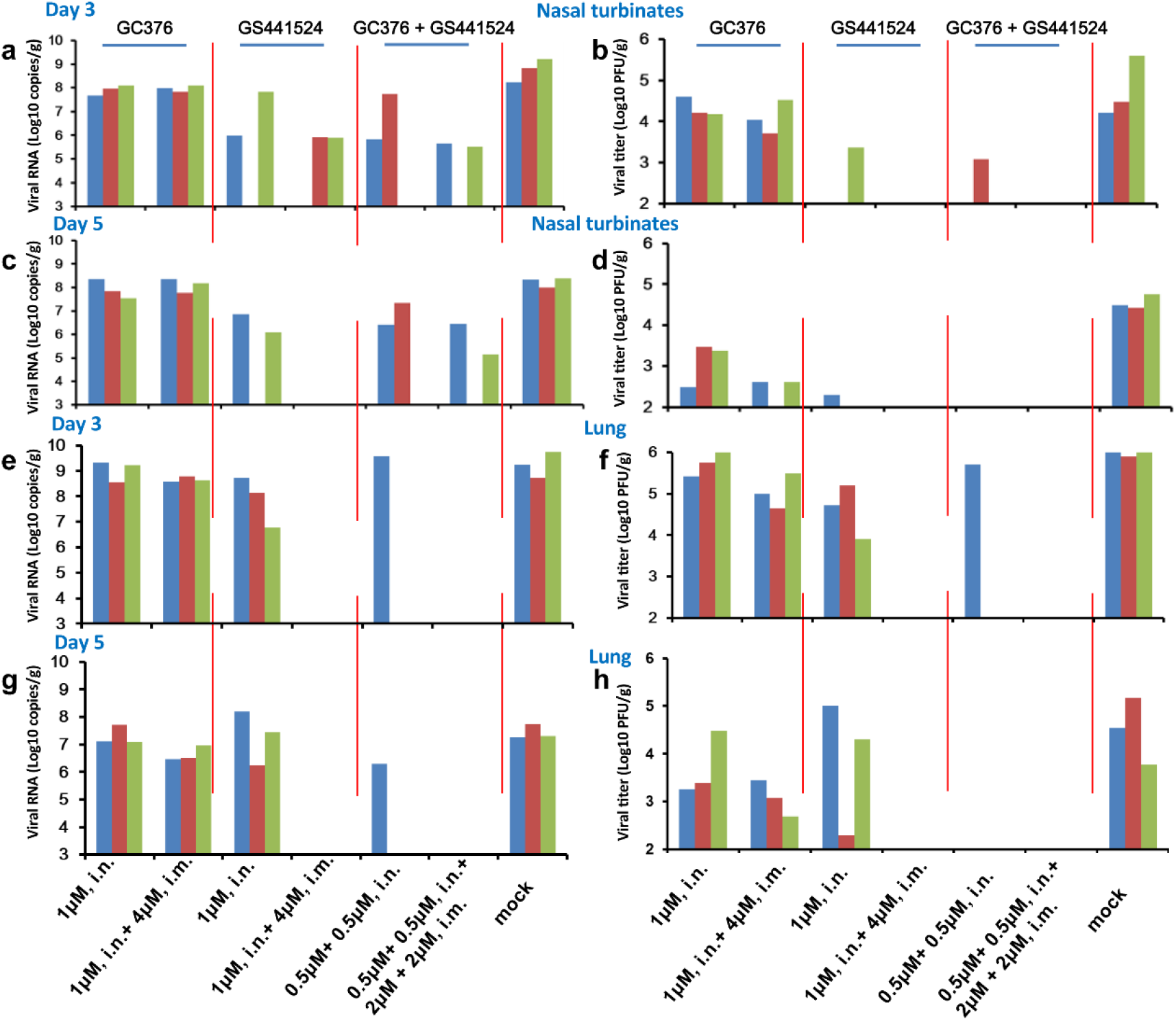
Evaluation of i.n. and i.m. GC376, GS441524 and GC376+GS441524 against SARS-CoV-2 infection in mice. Four- to six-week-old female BALB/c mice were administered a loading dose of GC376 (1 μM, i.n. or 1 μM, i.n.+4 μM, i.m.), GS441524 (1 μM, i.n. or 1 μM, i.n.+4 μM, i.m.), GC376 + GS441524 (0.5 μM+0.5 μM, i.n. or 0.5 μM+0.5 μM, i.n. and 2 μM+2 μM, i.m.), followed by a corresponding daily maintenance dose. Control mice were administered vehicle solution (12% sulfobutylether-β- cyclodextrin, pH 3.5) daily in parallel (0 μM). One hour after administration of the loading dose of GC376, GS441524, GC376 + GS441524 or vehicle solution, the mice were inoculated intranasally with 10^3.6^ PFU of HRB26M in a volume of 50 μl. On days 3 and 5 p.i., three mice in each group were euthanized and their nasal turbinates and lungs were collected. The viral RNA copies and infectious titers in the nasal turbinates (a-d) and lungs (e-h) were detected by qPCR and viral titration. The concentrations of the daily maintenance doses are shown.

### Pharmacokinetics study of GC376 and GS441524 alone or in combination

To further examine the potential of GC376 and GS441524, we evaluated their pharmacokinetic (PK) properties in SPF SD rats following i.m. administration of GC376 (111 mg/kg), GS441524 (67 mg/kg) and GC376+GS441524 (55.5+33.5 mg/kg), which are the same doses used in mice according to weight. The PK results showed that GC376 and GS441524 were rapidly absorbed after i.m. administration, and the peak plasma level was reached 1.30±0.60 h and 2.00±1.10 h after injection, respectively (**Fig. 5 and Table 1**). Our results are consistent with the previously reported T_max_ of GS441524 in patients (29) and GC376 in cats (18). We found that the maximum detected plasma drug concentration (C_max_) of GC376 (12.56±1.90 μg/ml) was approximately 2.5-fold lower than GS441524 (30.96±8.40 μg/ml) (**Table 1**). The value of the area under the curve (AUC_0-t_) of GS441524 (AUC_0-t_=183.33±64.36) was approximately 2.0-fold higher than GC376 (AUC_0-t_=92.14±9.99). However, the i.m. administered dose of GC376 was approximately 1.7-fold that of GS441524. We observed that the clearance rate of GC376 (CL/F, 1208±122 ml/h/kg) in plasma was approximately 2.9-fold that of GS441524 (CL/F, 423±186 ml/h/kg) (**Table 1**). These results indicated that the utilization efficiency of GS441524 *in vivo* is significantly higher than that of GC376. This finding may be one of the reasons for the poor ability of GC376 to inhibit SARS-CV-2 *in vivo*. In addition, current research has shown that GC376 can effectively eliminate FIPV in cats (18). However, this drug could not effectively inhibit SARS-CoV-2 in the upper and lower respiratory tracts in mice. One possible explanation is that GC376 may have difficulty reaching the mouse nasal turbinates and lungs, and the specific mechanism needs further study.

**Fig.5.**
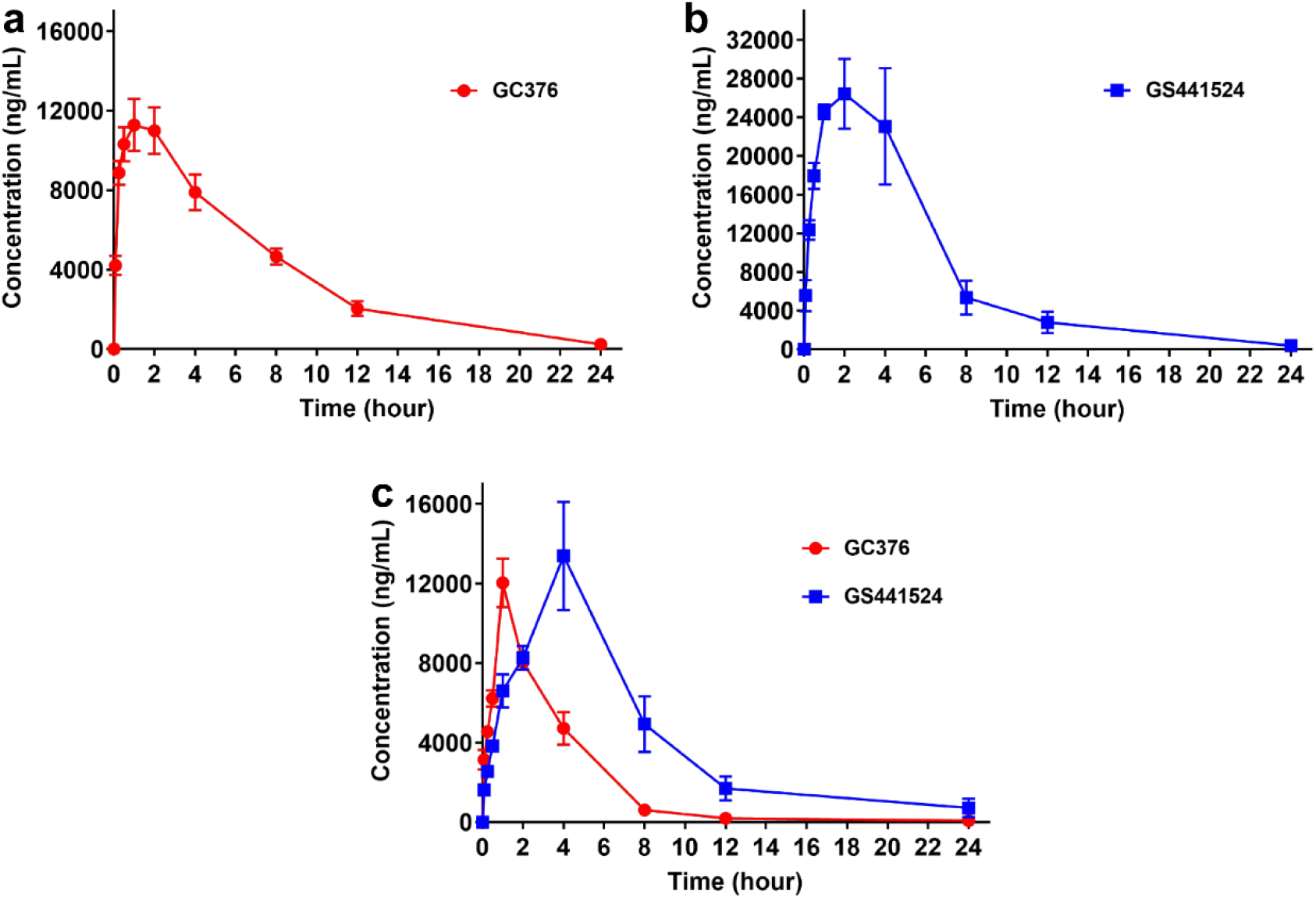
Plasma concentration-vs-time profiles following GC376, GS441524 and GC376+GS441524 administration in SD rats. In the single-dose PK study, five SPF SD rats were i.m. injected with GC376 (111 mg/kg), GS441524 (67 mg/kg) and GC376+GS441524 (55.5+33.5 mg/kg) for the determination of serial plasma drug concentrations. Data were analyzed via GraphPad Prism7.0, and the error bars show the SEM of the results from five replicates.

**Table 1.**
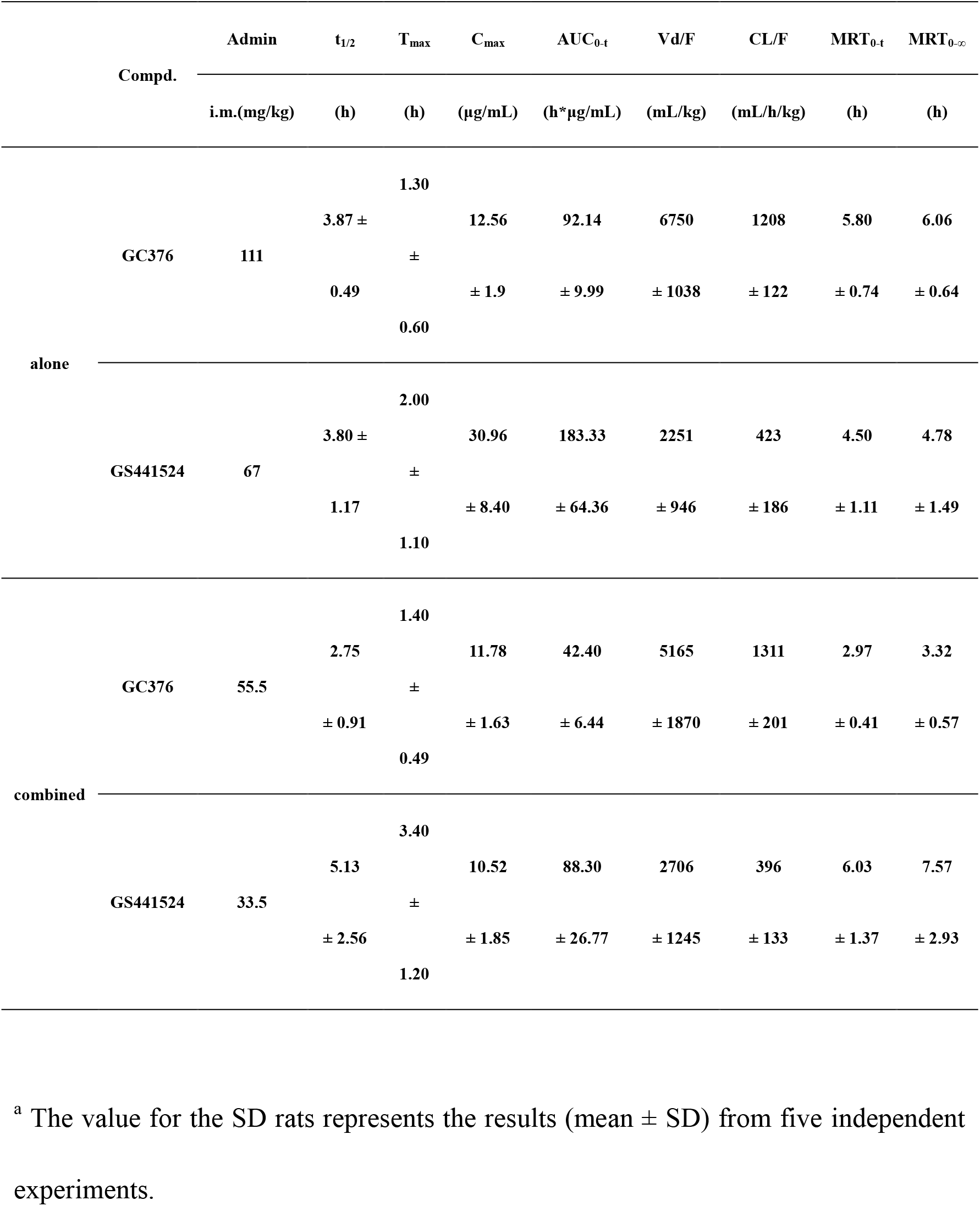
Preliminary pharmacokinetic (PK) evaluation of compounds GC376 and GS441524 in SD rats.^a^.

Furthermore, we found that the combined application of GC376 and GS441524 had a synergistic effect and extended T_1/2_ from 3.80±1.17 h to 5.13±2.56 h, T_max_ from 2.00±1.10 h to 3.40±1.20 h and the residence time of GS441524 (MRT_0-t_ from 4.50±1.11 h to 6.03±1.37 h) (**Fig. 5c and Table 1**). However, the injection dose was halved because the clearance rate was not changed, and the MRT_0-t_ of GC376 decreased from 5.80±0.74 h to 2.97±0.41 h. The PK study results showed that GC376 reached C_max_ earlier (T_max_=1.40±0.49) than GS441524 (T_max_=3.40±1.20 h) to produce a synergistic effect (**Fig. 5c)**. When these agents were combined, GC376 was the first drug to inhibit SARS-CoV-2 replication. After the plasma concentration of GC376 decreased, GS441524 reached its C_max_ (**Fig. 5c and Table1**) to produce a continuous inhibition of SARS-CoV-2 proliferation and maintain the effective concentration for a longer time. This phenomenon may explain why the combined application of GC376 and GS441524 was better than single application alone. In addition, the difference in the pharmacokinetics of GC376 and GS441524 between the upper and lower respiratory tract needs to be further evaluated.

In summary, we assessed the efficacy of GC376 and GS441524 to inhibit SARS-CoV-2 replication using a mouse-adapted virus infection model. Importantly, we found that intranasal administration of GS441524 and GC376+GS441524 significantly prevents the replication of virus in the upper respiratory tract, and the efficacy of GC376+GS441524 to inhibit the viral replication in the lower respiratory tract significantly better than that of GS441524. Combined i.n. and i.m. administration of GS441524 and GC376+GS441524 effectively protected mice against HRB26M infection in the upper and lower respiratory tracts, but GC376 alone failed to block the proliferation of SARS-CoV-2 in mice. Compared with GC376 and GS441524 alone, the dosage of GC376+GS441524 is halved, these results showed an additive effect of the combined application of the M^pro^ and RdRp inhibitors, so it should be developed and considered for future clinic practice.

## Materials and methods

### Compounds, cells, viruses and animal experiments

The compounds GC376 (molecular weight: 507.53 g/M) and GS441524 (molecular weight: 291.26 g/M) were synthesized at WuXi AppTec with purities higher than 95%. They were dissolved in 5% ethanol, 30% propylene glycol, 45% PEG 400, and 20% water with a concentration of 40 mM/l according to a previously described study (35).

Vero E6 cells (African green monkey kidney, ATCC) were maintained in Dulbecco’s modified Eagle’s medium (DMEM, Thermo Scientific, USA) containing 10% fetal bovine serum (Thermo Scientific, USA) and antibiotics, and incubated at 37°C with 5% CO2 (4). In addition, SARS-CoV-2/HRB26/human/2020/CHN (HRB26, GISAID access no. EPI_ISL_459909) and Mouse-adapted SARS-CoV-2/HRB26/human/2020/CHN (HRB26M, GISAID access no. EPI_ISL_459910) were obtained following the previous method (30). Infectious virus titers were determined using a plaque forming unit (PFU) assay in Vero E6 cells, and virus stocks were stored in aliquots at −80 °C until use.

Specific pathogen-free female BALB/c mice, aged 4-6 weeks were obtained from Beijing Vital River Laboratory Animal Technologies Co., Ltd (Beijing, China) and were housed and bred in the temperature-, humidity- and light cycle-controlled animal facility (20 ± 2 °C; 50 ± 10%; light, 7:00–19:00; dark, 19:00–7:00, respectively) of the Animal Center, Academy of Military Medical Sciences, Beijing.

### Protein expression and purification

SARS-CoV-2 M^pro^ flanked by an N-terminal His6 tag was cloned into pET-32a (+). For expression of SARS-CoV-2 M^pro^, the saved cells (from Sheng-ce Tao lab) were cultured at 37°C in LB medium containing 50 μg/ml ampicillin. Protein expression was induced with 0.8 mM isopropyl β-D-1-thiogalactopyranoside (IPTG) when the culture density reached an OD_600_ of 0.8, and cell growth continued for an additional 16 h at 18°C. Protein purification was performed as described previously (38, 39). Briefly, the cell supernatant was filtered with a 0.45-μm filter and loaded onto a nickel-charged HisTrap HP column (GE Healthcare). Proteins were eluted with elution buffer (20 mM Tris-HCl, 500 mM NaCl and 500 mM imidazole, pH 7.4). Then, the harvested protein was concentrated to approximately 2.0 ml and filtered using a Superdex 200 gel filtration column (GE Healthcare) equilibrated with buffer (20 mM Tris-HCl and 200 mM NaCl, pH 7.4). For crystallization, the purified protein was concentrated to approximately 8 mg/ml, flash frozen with liquid nitrogen, and stored at −80°C. The concentration of purified SARS-CoV-2 M^pro^ was determined by the absorbance at 280 nm (A_280_) using a NanoDrop 2000c UV-Vis spectrophotometer (Thermo Fisher Scientific).

### Crystallization, data collection and structure determination

GC376 and M^pro^ (8 mg/ml) were incubated at room temperature for 1 h (molar ratio: 1:2), and the complex was crystallized by the hanging drop vapor diffusion method at 20°C. The best crystallization conditions for the complex were in hanging drops consisting of 2 μl of reservoir solution (0.1 M sodium malonate pH 6.0, 12% w/v polyethylene glycol 3,350) and 2 μl of the complex in 20 mM Tris-HCl and 200 mM NaCl, pH 7.4, followed by incubation at 20°C for 3 days. Then, the crystals were flash-cooled in liquid nitrogen in a cryoprotectant solution containing 30% ethylene glycol and 70% reservoir solution (0.1 M sodium malonate pH 6.0, 14% w/v polyethylene glycol 3,350). Data collection was performed at the research associates at Center for Protein Research (CPR), Huazhong Agricultural University (wavelength = 1.5418 Å, temperature = 100 K). Reflections were integrated, merged, and scaled using HKL-3000 (40), and the resulting statistics are listed in **Table S1**. The structure was solved by molecular replacement with PHASER (Phenix, Berkeley, CA, USA) (41) using the structure of the SARS-CoV-2 M^pro^ (PDB identifier 7BQY) as a starting model. Manual model rebuilding and refinement were performed using COOT (42) and the PHENIX software suite. Structural figures were generated using PyMOL (Schrödinger). Detailed molecular interactions were determined using LIGPLOT (43).

### Evaluation of antiviral activity in Vero E6 cells

Cell viability was determined using the Cell Titer-Glo kit (Promega, Madison, WI, USA) following the manufacturer’s instructions. Briefly, Vero E6 cells were seeded in 96-well plates with opaque walls. After 12 to 16 h, the indicated concentrations of GC376 (0, 1, 5, 10, 50, 100, 500 μM), GS441524 (0, 1, 5, 10, 50, 100, 500 μM) and GC376+GS441524 (0, 0.5, 2.5, 5, 25, 50, 250μM) were added for 24h. Cell Titer-Glo reagent was added to each well, and luminescence was measured using a GloMax 96 Microplate Luminometer (Promega, Madison, WI, USA).

Antiviral activity experiment was determined following a previous method (30). Briefly, Vero E6 cells were pretreated with the indicated concentrations of GC376 (0, 0.5, 1, 2, 4, 6, 8, 10 μM), GS441524 (0, 0.5, 1, 2, 4, 6, 8, 10 μM) and GC376+GS441524 (0, 0.25, 0.5, 1, 2, 3, 4, 5 μM) or with vehicle solution (12% sulfobutylether-β-cyclodextrin, pH 3.5) alone for 1 h. The cells were then infected with HRB26 or HRB26M at an MOI of 0.005 and incubated for 1 h at 37°C. The cells were washed with PBS, and virus growth medium containing the indicated amounts of GC376, GS441524 and GC376+GS441524 or vehicle solution alone was added. The supernatants were collected at 24 h p.i. for viral titration by a PFU assay in Vero E6 cells. Relative viral titers were calculated on the basis of the ratios to the viral titers in the mock-treated counterparts. The data were analyzed using GraphPad Prism 7.0. The results are shown as the mean values with standard deviations of three independent experiments.

### *In vivo* toxicity study of GC376 and GS441524

The toxicity studies were performed in 4- to 6-week-old female BALB/c mice. BALB/c mice were assigned to four groups (five mice per group), one mock group (i.m. administration of solvent) and three i.m. administered groups: GC376 (40 mM/l, 100 μl), GS441524 (40 mM/l, 100 μl) and GC376+GS441524 (20 mM/l, 100 μl), respectively. Mice in the mock and experimental groups were weighed daily for 15 days. In addition, blood samples were collected at 0, 5, 10 and 15 days after administration. Various blood chemistry values or blood cell counts were performed at Wuhan Servicebio Biological Technology Co., Ltd. The data were analyzed using GraphPad Prism 7.0.

### *In vivo* antiviral study of GC376 and GS441524

Firstly, groups of six 4- to 6-week-old female mice were treated i.m. with a loading dose of GC376 (40 or 8 mM/l, 100 μl), GS441524 (40 or 8 mM/l, 100 μl) and GC376+GS441524 (20 or 4 mM/l, 100 μl), followed by a daily maintenance dose. Alternatively, mice were treated intranasally with a single treatment (GC376, 20 mM/l, 50 μl; GS441524, 20 mM/l, 50 μl; GC376+GS441524, 10 mM/l, 50 μl) or a combination of GC376 (20 mM/l, 50 μl, i.n. and 40 mM/l, 100 μl, i.m.), GS441524 (20 mM/l, 50 μl, i.n. and 40 mM/l, 100 μl, i.m.) and GC376+GS441524 (10 mM/l, 50μl, i.n. and 20 mM/l, 100 μl, i.m.), followed by a daily maintenance dose. As a control, mice were administered vehicle solution (12% sulfobutylether-β-cyclodextrin, pH 3.5) daily. One hour after administration of the loading dose of GC376, GS441524 and GC376+GS441524 or vehicle solution, each mouse was inoculated intranasally with10^3.6^ PFU of HRB26M in 50 μl. Three mice from each group were euthanized on days 3 and 5 p.i. The nasal turbinates and lungs were collected for viral detection by qPCR and PFU assay according to previously described methods (4, 30). The amount of vRNA for the target SARS-CoV-2 N gene was normalized to the standard curve from a plasmid (pBluescript II SK-N, 4,221 bp) containing the full-length cDNA of the SARS-CoV-2 N gene. The assay sensitivity was 1000 copies/ml. The data were analyzed using Microsoft Excel 2016 and GraphPad Prism 7.0.

### Pharmacokinetics study of GC376 and GS441524 in SD rats

Five healthy SPF SD rats of 4-6 weeks were used in a single-dose PK study. At time point zero, the SD rats of groups A, B and C received i.m. injections of GC376 (111 mg/kg), GS441524 (67 mg/kg) and GC376+GS441524 (55.5+33.5 mg/kg), which are the same doses used in mice according to weight. Approximately 200μl of blood was collected at 0, 0.083, 0.25, 0.5, 1, 2, 4, 8, 12, and 24 h from the tail vein and placed in a precooled polypropylene centrifuge tube containing 3.0 μl of 40% EDTAK2. Then, the whole blood was centrifuged at 7800 g/min for 10 min at 4℃. Plasma was collected and stored in a freezer at −80°C. Plasma drug concentration was analyzed using LC-MS/MS. Pharmacokinetic parameters were calculated using WinNonlin software (version 6.4), and a non-atrioventricular model was used for data fitting. The data were analyzed using Microsoft Excel 2016 and GraphPad Prism 7.0.

### Ethics statement, biosafety and facility

All of the mice used in this study were maintained in compliance with the recommendations in the Regulations for the Administration of Affairs Concerning Experimental Animals made by the Ministry of Science and Technology of China. The toxicity studies in mice and single-dose PK study in SD rats were performed using protocols that were approved by the Scientific Ethics Committee of Huazhong Agricultural University. All experiments with infectious SARS-CoV-2 were performed in the biosafety level 4 and animal biosafety level 4 facilities in the Harbin Veterinary Research Institute (HVRI) of the Chinese Academy of Agricultural Sciences (CAAS), which are approved for this use by the Ministry of Agriculture and Rural Affairs of China.

### Statistical analysis

Statistical analysis was carried out using GraphPad Prism 7.0. Statistical significance was determined using an unpaired two-tailed Student’s t test. Data are presented as mean values +/− SD (95% confidence interval). *, P <0.05 was considered statistically significant; **, P < 0.01 was considered highly significant; ***, P < 0.001 and ****, P < 0.0001 were considered extremely significant. All experiments were further confirmed using biological repeats.

## Data availability

Coordinates and structural factors for SARS-CoV-2 M^pro^ in complex with GC376 were deposited in the Protein Data Bank with accession number 7CBT. Other data supporting the findings of this study are available within the paper and its Supplementary Information files, or upon reasonable request from the corresponding author.

## Acknowledgments

We thank research associates at the Center for Protein Research (CPR), Huazhong Agricultural University, for technical support. We also thank Professor Sheng-ce Tao from Shanghai Jiao Tong University for providing SARS-CoV-2 M^pro^ plasmid. This work was supported by National Natural Science Foundation of China Grants 31722056, the National Key R&D Program of China (Grant No. 2018YFC1200601), the Natural Science Foundation of Hubei Province of China (Grant No.2020FCA040), the Fundamental Research Funds for the Central Universities (Grant No.2662020PY001) and the Applied Technology Research and Development Project of Heilongjiang Province, China (Grant No. GA20C006).

## Author contributions

G.Q.P., Z.G.B., Z.F.F. and H.C.C. designed the study; Y.J.S., L.S., Z.Y.W., C.W., Y.Y.Y, Z.J. and F.L.G. performed the experiments; G.Q.P. and Z.G.B. oversaw the project; Y.J.S., L.S., Z.F.F., H.C.C., Z.G.B and G.Q.P. analyzed the data and wrote the manuscript.

## Competing interests

The authors declare no competing interests.

## Additional information

Correspondence and requests for materials should be addressed to G.Q.P. or Z.G.B..

